# The Effects of Acute Trazodone Administration on Sleep in Mice

**DOI:** 10.1101/2025.01.22.634422

**Authors:** Mayuko Arai, Brianne A Kent

## Abstract

**Study Objectives:** Trazodone is an antidepressant with robust hypnotic effects, frequently prescribed off-label to treat insomnia. Trazodone has gained recent attention in the context of neurodegenerative diseases because sleep has been proposed as a novel target for disease-modifying therapeutics. Preclinical research in rodents examining the effects of trazodone on sleep is limited, so here we aimed to develop a translationally-focused protocol to study the sleep-promoting effects of trazodone in mice.

**Methods:** We investigated the effects of voluntary oral trazodone administration at doses of 0 mg/kg, 10 mg/kg, 40 mg/kg, and 60 mg/kg on sleep in C57BL/6J mice (n = 15; females = 6; age 10-13 months). Mice were dosed with trazodone for 6 consecutive nights, while being recorded with intracranially implanted 2-channel electroencephalogram (EEG) and electromyography (EMG). EEG/EMG recordings were analyzed for time spent in each vigilance state and power spectra.

**Results:** A single dose of trazodone, administered prior to the onset of the 12-h rest phase, dose-dependently increased non-rapid eye movement (NREM) sleep and delta power during NREM sleep, at the expense of rapid eye movement (REM) sleep. These effects on sleep persisted after six consecutive days of dosing, albeit to a lesser extent.

**Conclusion:** We have validated a novel voluntary oral administration protocol for trazodone use in mice and have shown that trazodone effectively promotes NREM in mice. Our novel protocol will be useful for future research investigating the effects of trazodone on sleep in mouse models of disease.

**Statement of significance:** Trazodone is an antidepressant known to increase slow wave sleep in humans. Slow wave sleep promotion is being explored as a disease-modifying therapeutic target for neurodegenerative diseases, such as Alzheimer’s disease. We have developed a novel translationally-focused protocol for administering trazodone to mice, which overcomes some of the challenges of using trazodone for preclinical sleep research. Using our protocol we have demonstrated for the first time that trazodone dose-dependently increases the amount of NREM sleep and delta power in mice, recapitulating the enhancement of slow wave sleep in humans treated with trazodone.

## Introduction

Trazodone is an anti-depressant, primarily acting as a serotonin 5-HT2A antagonist and serotonin reuptake transporter (SERT) inhibitor [1]. In addition to the anti-depressant effects, trazodone has robust hypnotic actions and is commonly prescribed for off-label use to treat sleep disorders at a lower dosage (25-100 mg) than when prescribed to treat depression (150-600mg) [1,2]. A recent observational study in long-term care facilities found that about 50% of the older adults treated with trazodone were prescribed the medication for agitation or insomnia, with or without a diagnosis of depression [3].

Trazodone specifically increases the amount of slow wave sleep (SWS), defined by low frequency EEG oscillations occurring during the N3 stage of NREM sleep [4]. The sleep-promoting effects of trazodone, particularly the enhancement of SWS, were first observed by Montgomery and colleagues (1983) in elderly participants [4]. Their study found that while trazodone did not extend total sleep duration, trazodone significantly increased the duration of SWS. Subsequent research has replicated these findings, demonstrating increases in SWS or NREM sleep more generally [5–12].

Pharmacological agents that can increase SWS are gaining attention in the context of neurodegenerative diseases, because SWS is a critical aspect of sleep regulating perivascular convective cerebrospinal fluid (CSF) flow in the brain [13,14]. Interest in trazodone as a potential therapy for disorders that are associated with reduced SWS, such as Alzheimer’s disease, has also grown recently due to its ability to alleviate sleep disturbances in the elderly with minimal adverse effects [15–17]. Notably, research from the UCSF Memory and Aging Center found that trazodone was associated with slower cognitive decline in elderly users [18], and a randomized controlled trial is currently underway to evaluate its effects on cognitive decline in older adults diagnosed with amnestic mild cognitive impairment with sleep complaints [17].

Despite the growing interest in trazodone, preclinical research beyond the antidepressant properties remains limited. Previous rodent studies have struggled to replicate the sustained SWS-promoting effects observed in humans. These inconsistencies may stem from issues related to dosage and administration methods [19,20]. Specific challenges for administering trazodone to rodents include: (1) trazodone dissolved in solution is too acidic to safely administer via injection (without changing the composition of the solution) and (2) administering trazodone via oral gavage is not possible with the design of most intracranial EEG/EMG implants needed for assessing sleep architecture. Establishing a reliable method for studying the effects of trazodone on sleep in rodents is crucial for deepening our understanding of trazodone as a sleep aid and for exploring its therapeutic potential in sleep-related disorders. In this study, we validated a novel voluntary oral administration protocol that can be used to characterize the effects of trazodone on sleep architecture in mice.

## Method

### Animals

All experimental protocols were approved by the Simon Fraser University Animal Care and Use Committee (Protocol #1365P-23). A total of 15 (female n = 6) C57BL6/J mice (age 10-13 months, Charles River Laboratories, Quebec, Canada) were used and single-housed under a 12:12h light/dark cycle after the implantation surgery.

### EEG/EMG implantation and recordings

The animals were implanted with a 2-channel EEG/1-channel EMG headmount (#8201-SS; Pinnacle Technology, Lawrence, KS, United States). Throughout the EEG implantation procedure, the animals were anesthetized deeply with isoflurane. The cranial implant consisted of four stainless steel screws at the following coordinates relative to bregma: AP: +/− 3 mm, ML: +/− 1.5 mm. The screw electrodes were inserted through holes of the prefabricated EEG headmount and rotated into 4 burr holes drilled through the skull. Two EMG electrode wires attached to the headmount were inserted under the nuchal muscles. A layer of dental cement (A-M Systems) was applied to secure the headmount to the skull. Meloxicam (5mg/kg, IP), buprenorphine (0.07mg/kg, SC), and lidocaine (7mg/kg, SC along the incision site) were administered for analgesia at the beginning of the surgery. Lactated Ringer’s (10mg/kg) was administered at the end of surgery to prevent dehydration. Meloxicam and buprenorphine were also provided for 2 days post-op. All mice were given at least 7 days to recover before EEG/EMG recordings.

For the EEG/EMG recordings, the mice were habituated for at least 24 h to a rectangle clear plexiglass housing cage used for the recordings and the wireless EEG unit (#8274-SL; Pinnacle Technology). After the habituation period, all animals were recorded with an EEG monitoring system (#8200-K9-SL; Pinnacle Technology) for two days as a baseline and then 6 days while receiving the trazodone treatment or vehicle. The recording chamber was maintained on a regular 12:12 LD cycle with *ad libitum* access to water. The EEG signals were recorded at a sampling rate of 256 Hz, with a total gain of 2600 V/V. A high-pass filter was set at 0.5 Hz for EEG recordings and 10 Hz for EMG recordings.

### Trazodone administration

Trazodone hydrochloride (HCl) powder (catalog number T6154; Sigma-Aldrich) was dissolved in distilled water to achieve 10-50mg/ml concentration, and then mixed with palatable food, immediately before the daily administration. The concentration was adjusted to maintain a consistent moisture level in the food mixture and to achieve one of the four doses (0mg as the control; 10mg/kg, 40mg/kg, and 60mg/kg as the treatments), without altering the total volume of the dissolved trazodone solution (0.8– 1.2 ml/kg). Trazodone is acidic and not all palatable foods mask the taste enough to encourage consumption by mice. After piloting several types of food mixtures (e.g., peanut butter, see Supplementary Materials, Table S1 for a complete list of foods that were piloted), the palatable food selected for the present study was a strawberry-flavored treat (VitaKraft® Drops with Strawberry, Product #25451, Vitakraft Sunseed, Bowling Green, OH, United States). Following the one-week recovery period after the EEG/EMG implantation surgery, the amount of daily rodent food pellets was reduced gradually, aiming for 90-95% of the baseline body weight. The food pellets were given daily at ZT12.5 (0.5 h after lights-off, which is the start of the active phase in nocturnal rodents). From day 3 of food restriction, animals were introduced to the palatable food without trazodone daily at ZT 23 (1 h prior to lights-on, which is the start of the rest phase). Once the animals began ingesting the treat within 15 minutes, they were transferred to the EEG recording cage and habituated for at least 24 hours. Then, each animal was provided with palatable food at ZT 23.5 for two days to establish a baseline level of sleep. Following this, trazodone treatment was administered at ZT 23.5 for six consecutive days. Dosing was calculated based on daily body weight, recorded at ZT23.

After the six-day dosing period, the animals were returned to their regular housing cages for a washout period of at least three weeks. During the washout phase, palatable food was given without trazodone every five days to maintain familiarity with the palatable food. Following the washout period, each animal was recorded for two baseline days to confirm the washout’s completion. Subsequently, they received a second 6-day dose of trazodone. The doses were counterbalanced, so that animals receiving the highest dose initially were given the lowest dose in the second round, and vice versa.

### Sleep Analysis

For vigilance state and power spectra analyses, EEG traces were scored for wake, NREM, and REM in 10-s epoch durations using Sirenia Sleep Pro software (Pinnacle Technology) by an investigator blind to the treatment condition. Sleep scoring of the EEG recordings and power spectral analyses were conducted following the methods described previously [21,22]. Briefly, the data were first cluster scored by grouping epochs based on EEG and EMG frequency bands (e.g., delta, theta, alpha, beta, gamma), identifying bouts of NREM, REM and wake. The epoch classifications were then visually confirmed by an experimenter reviewing EEG/EMG, video recordings, and corresponding spectral plots. Wake was defined by low-amplitude EEG (dominant frequency above 4 Hz) and high-amplitude EMG. NREM sleep was defined by high-amplitude EEG and peak frequencies below 4 Hz, and low-amplitude EMG. REM was defined by EEG peak frequencies between 4 and 8 Hz, uniform low amplitude EEG waveforms, low-amplitude EMG, and transitioning from NREM. Epochs were categorized based on the predominant state (>50%) of each epoch. To analyze the power spectrum, the Fast Fourier Transform was used. The frequency bands were defined as delta (1–4 Hz), theta (4–8 Hz), alpha (8–12 Hz), beta (12–30 Hz), and gamma (30–50 Hz), and a bandpass filter with a range of 1-50 Hz was applied to all the data.

### Data analysis

Power spectra were analyzed as a percentage of total power across the frequency range of 1 Hz to 50 Hz. Statistical analyses were conducted using R (version 4.3.3). To evaluate the effects of trazodone on the duration of vigilance states across different doses, we performed mixed-design ANOVAs, followed by post hoc Tukey HSD tests when there were statistically significant interactions. To compare power between baseline and treatment days within each dose, we used repeated-measures ANOVAs with post hoc Tukey HSD tests. When the assumption of sphericity was violated, the Greenhouse-Geisser correction was applied. Additionally, to compare power across doses for each frequency band, we conducted two-way ANOVAs (dose × EEG electrode), followed by Tukey HSD post hoc tests. All data are presented as mean ± standard error of the mean (SEM).

## Results

### Mice voluntarily consume trazodone mixed with palatable food

When mice were initially introduced to the palatable food, it took an average of 3.4 days (SD ±1.82) to consume the palatable food within 15 min. One male mouse stopped consuming the palatable food during the baseline recording and was excluded from the study. The remaining 14 mice proceeded with the trazodone dosing. 12 of the 14 mice received 2 separate doses of trazodone with a 3-week washout period between doses. 2 of the 14 mice only received 1 dose of trazodone and were not retested due to poor EEG quality by the second testing timepoint (the EEG/EMG headmount can become loose over time and cause excessive artifact in the EEG signal). The final sample sizes for each dose were: n=2 vehicle 0mg/kg, n=8 10mg/kg trazodone, n=8 40mg/kg trazodone, and n=8 60mg/kg trazodone. On Day 1 of dosing, all mice consumed the entire vehicle or treatment dose. On Day 6 of dosing, n=2 from the 40mg/kg condition and n=2 from the 60mg/kg condition did not consume the full dose and were exuded from the Day 6 analyses.

### A single dose of trazodone dose-dependently increases NREM sleep duration

Following the first dose of trazodone, we compared time spent in each vigilance state to baseline values (average of two baseline days) during the 12-h lights-on phase (i.e., rest phase). Four mice were excluded from these analyses due to equipment issues, specifically, the wireless EEG disconnected during the recording or had poor signal quality. Trazodone administration (0, 10, 40, and 60 mg/kg) had a dose-dependent effect on the duration of NREM, REM, and wake (Figure 1); mixed-design ANOVA revealed a significant interaction between the trazodone dose and the duration of vigilance state (F(6, 36) = 17.89, p < 0.0001). Post-hoc Tukey tests showed that compared to the 0 mg/kg control, significant increases in NREM duration were observed at the 10 mg/kg (t(54) = 3.96, p = 0.0012), 40 mg/kg (t(54) = 6.52, p < 0.0001), and 60 mg/kg doses (t(53.4) = 8.80, p < 0.0001). Additionally, NREM duration increased progressively with dose: significant differences were found between 60 mg/kg and 40 mg/kg (t(50.1) = 3.38, p = 0.0074), 60 mg/kg and 10 mg/kg (t(52.8) = 6.82, p < 0.0001), and 40 mg/kg and 10 mg/kg (t(52.8) = 3.58, p = 0.0040) (Fig. 1A). For REM sleep, duration decreased at higher doses (e.g., 60 mg vs. 10 mg, t(52.8) = - 4.89, p < 0.0001; 40 mg vs. 10 mg, t(52.8) = - 4.13, p = 0.0007) (Fig. 1B). Wake duration also decreased across doses, with significant reductions compared to the control (e.g., 60 mg/kg vs. 0 mg/kg, t(53.4) = - 4.81, p = 0.0001) (Fig. 1C). The increase in NREM was also seen when assessing the entire day (22 h) post-administration (see Supplementary Materials, Figure S1). The data separated by sex are provided in the supplementary materials, but our study was underpowered to statistically analyze the effects of sex (Supplementary Figure S2).

**Figure 1.**
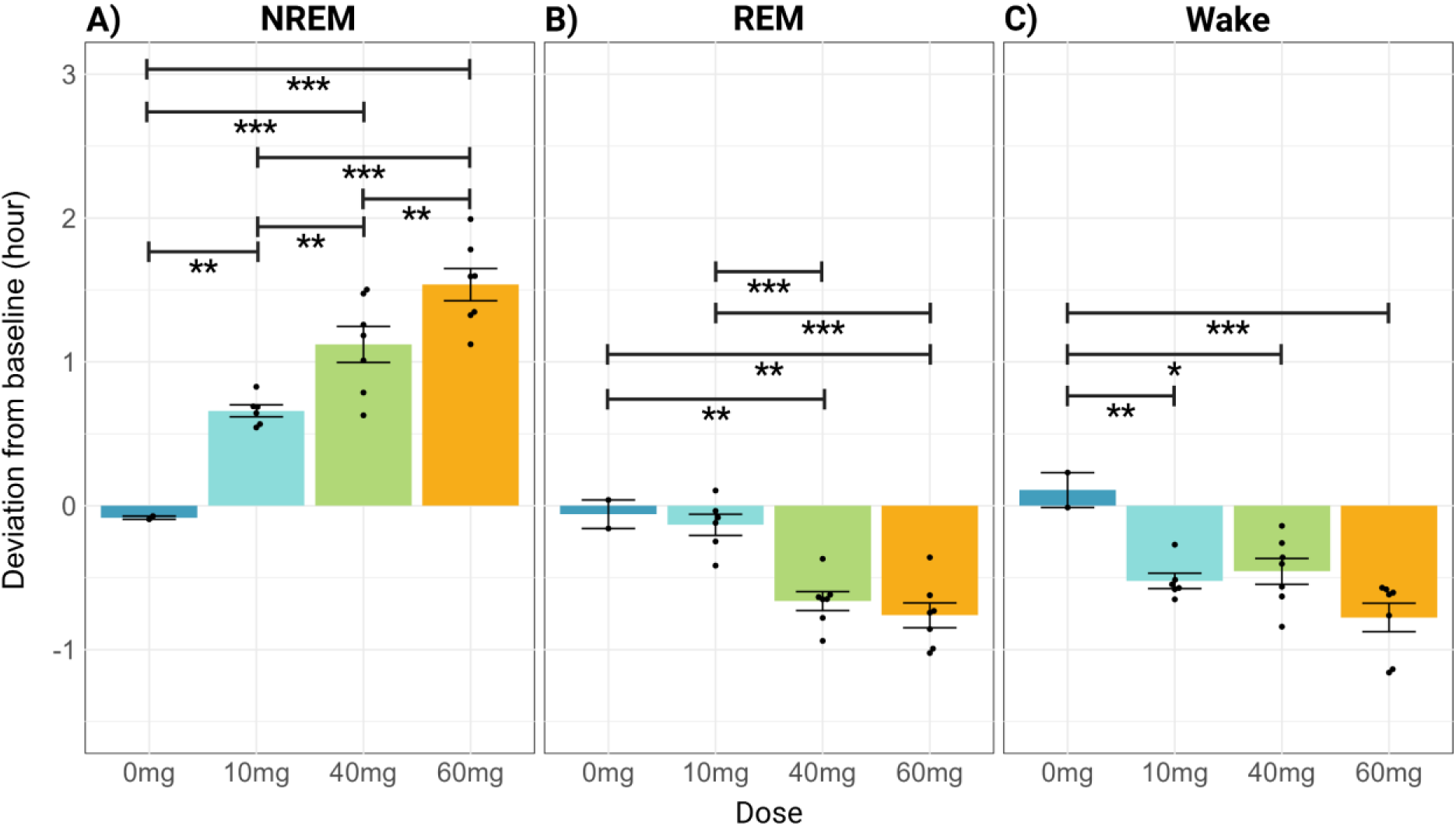
Acute trazodone administration dose-dependently affects the duration of vigilance states. The difference in (A) NREM, (B) REM, and (C) wake duration between Day 1 of trazodone administration and baseline, expressed in hours along the y-axis, for mice treated with trazodone HCL mixed with palatable food. The x-axes show the dose administered. Data are expressed as the mean ± SEM. Mixed design ANOVA followed by Tukey’s HSD test ∗ p < 0.05, ∗∗p < 0.01, ∗∗∗p < 0.001.

### A single dose of trazodone dose-dependently increases delta power

We compared the EEG power spectra following the first dose of trazodone to baseline levels (Figure 2). Power was normalized as a proportion of the total power within each vigilance state. Five mice were excluded from this analysis due to the wireless EEG disconnecting or exhibiting poor signal quality. At the 60 mg/kg dose, during NREM sleep (Fig. 2A), repeated-measures ANOVA revealed statistically significant interactions between conditions (baseline vs. treatment) and frequency bands at the frontal electrode (F(4, 24) = 25.92, p < 0.001). Post hoc Tukey HSD analysis showed that the 60mg/kg dose was associated with a significant increase in delta power (t(6) = 6.76, p = 0.0077) accompanied by a reduction in alpha (t(6) = - 8.34, p = 0.0025) and beta (t(6) = - 5.24, p = 0.0273) power. Similarly, at the 40 mg/kg dose (Fig. 2D), a significant interaction between the treatment condition and frequency band was observed at the frontal electrode (F(4, 20) = 18.90, p < 0.001) during NREM sleep. Post hoc analyses indicated a significant reduction in alpha power (t(5) = - 7.08, p = 0.0111) followed the 40mg/kg dose, but changes in delta power were not statistically significant (t(5) = 3.93, p = 0.1173). In contrast, no significant interactions were observed at the 10 mg/kg dose (Fig. 2G).

**Figure 2.**
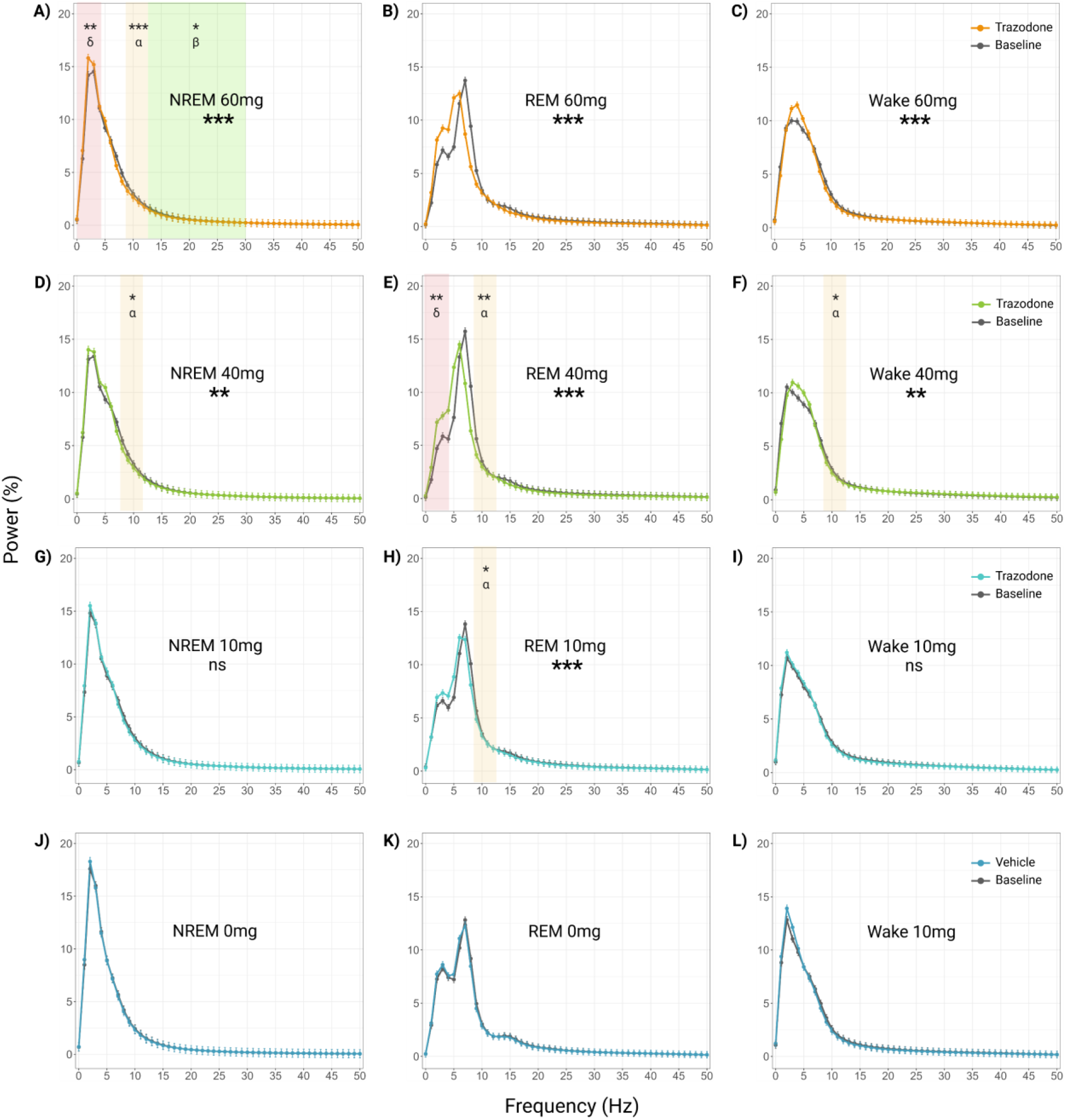
Acute trazodone administration dose-dependently shifts EEG power spectra. EEG power spectra recorded by the frontal electrode during NREM, REM, and wake following a single trazodone dose of 60mg/kg (A-C), 40mg/kg (D-F), 10mg/kg (G-I), and 0mg/kg (J-L). Data are expressed as the mean ± SEM. Repeated-measures ANOVA followed by Tukey’s HSD test. ^∗^ p < 0.05, ^∗∗^p < 0.01, ^∗∗∗^p < 0.001. δ = delta frequency, α = alpha frequency, β = beta frequency.

For REM sleep, significant interactions between the treatment condition and frequency band were observed at both the 60 mg/kg (F(4, 16) = 18.80, p < 0.001; note that two animals did not spend time in REM) and 40 mg/kg doses (F(4, 20) = 40.54, p < 0.001) in the frontal electrodes (Fig. 2B, 2E). Post hoc analysis revealed a significant increase in delta power at the 40 mg/kg dose (t(5) = 9.12, p = 0.0035) and a reduction in alpha power at the same dose (t(5) = -8.03, p = 0.0063). At the 10 mg/kg dose, a significant interaction was detected in the frontal electrode (F(2.09, 10.45) = 9.83, p = 0.004) when the Greenhouse-Geisser correction was applied (Fig. 2H).

During the wake period, significant interaction effects in the frontal electrode were observed at the 60 mg/kg dose (F(4, 24) = 9.01, p = 0.0001) and at the 40 mg/kg dose with Greenhouse-Geisser correction applied (F(4, 24) = 9.01, p = 0.044) (Fig. 2C, 2F). Post hoc analysis using Tukey HSD correction noted a significant reduction in the alpha band at the 40 mg/kg dose (t(5) = -6.92, p = 0.0122). No significant interaction was found at the 10 mg/kg dose (Fig. 2I).

Statistical analysis was deferred for the 0mg group due to the small sample size (n=2); however, Figures 2J – 2L illustrate the overlap between the baseline and post-vehicle administration for each vigilance state, confirming no effect from the vehicle control.

Similar results were observed in the power spectra from EEG recorded from the parietal electrodes across doses; data provided in the Supplementary Materials (Supplementary Fig. S3).

To assess the impact of trazodone on EEG spectra power across different doses, we analyzed deviations in power from baseline for each frequency band (Figure 3). Five animals were excluded from this analysis due to wireless EEG malfunctions or poor signal quality. Additionally, three other mice were excluded only from the parietal EEG electrode analysis due to the high levels of artifact in the EEG signal. For the delta band (defined as 1–4 Hz, Fig. 3A), a two-way ANOVA revealed a significant main effect of dose on power (F(3,31) = 5.477, p = 0.0039), while the interaction between dose and EEG electrode position was not statistically significant (F(3,31) = 0.420, p = 0.7402), suggesting that the dose effect was consistent across electrode positions. Post hoc comparisons indicated a significantly greater delta enhancement at the 60 mg/kg dose compared to the 10 mg/kg dose (t(31) = 3.09, p = 0.0210) and the 0 mg/kg control (t(31) = 3.40, p = 0.0096). This increase in delta power was accompanied by a reduction in alpha power (defined as 8–12 Hz, Fig. 3C), with a significant main effect of dose (F(3,31) = 12.272, p < 0.0001). Post hoc tests revealed stronger reductions in alpha power at higher doses, particularly at the 60 mg/kg dose compared to the 10 mg/kg dose (t(31) = - 4.10, p = 0.0015) and the 0 mg/kg control (t(31)= - 5.49, p < 0.0001). Additionally, a significant effect of dose was also observed in the theta frequency band (defined as 4–8 Hz, Fig. 3B) (F(3, 31) = 3.533, p = 0.0261) while the post hoc analysis did not identify any specific pairwise comparisons with statistically significant differences. No significant effects of dose or dose-by-electrode interactions were observed for the beta or gamma bands (Figs. 3D, 3E).

**Figure 3.**
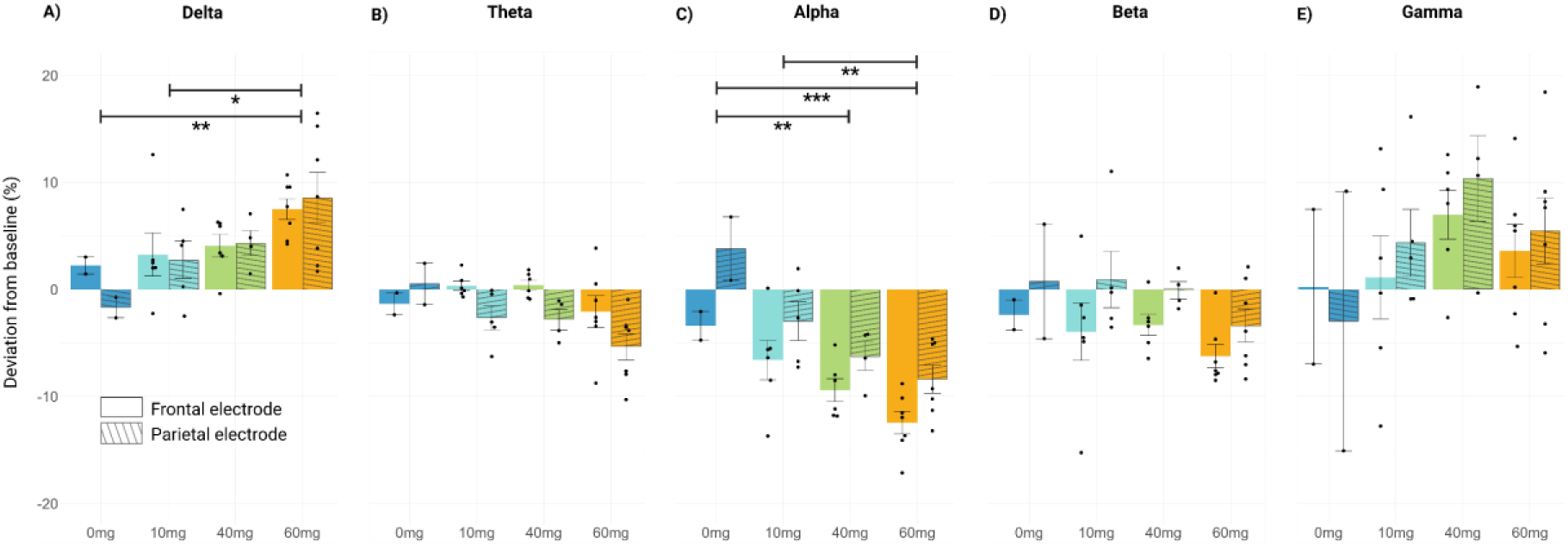
A single dose of trazodone increases delta power during NREM sleep, dose-dependently. The percent change (y-axis) from baseline in EEG spectra power for (A) delta, (B) theta, (C) alpha, (D) beta, and (E) gamma frequency bands during NREM sleep. X-axes show trazodone dose administered (0mg/kg, 10mg/kg, 40mg/kg, 60mg/kg). Data from the frontal electrode are represented as bars solid colours and data from the parietal electrode are represented by bars with diagonal lines. Data are expressed as the mean ± SEM. Two-way ANOVA followed by Tukey’s HSD test. ∗ p < 0.05, ∗∗p < 0.01, ∗∗∗p < 0.001.

### Trazodone maintains increased EEG delta spectra power on day 6 of administration

On Day 6 of administration, trazodone continued to exhibit a NREM-promoting effect (Fig. 4A), although the differences from baseline were less pronounced compared to Day 1. Five animals were excluded from this analysis due to the wireless EEG disconnecting during the recording or poor EEG quality. A mixed-design ANOVA indicated an interaction between dose and vigilance state was not significant (F(6, 26) = 0.633, p = 0.149), after applying the Greenhouse-Geisser correction. However, NREM sleep was still enhanced compared to baseline, increasing by 0.78 hours on average (± 0.40 hours SEM) at the 60 mg dose. When evaluating the entire day (22 hours), an increase in NREM sleep duration was not observed on Day 6 (Supplementary Materials Figure S1). The data separated by sex is provided in the Supplementary Materials (Supplementary Figure 2B).

**Figure 4.**
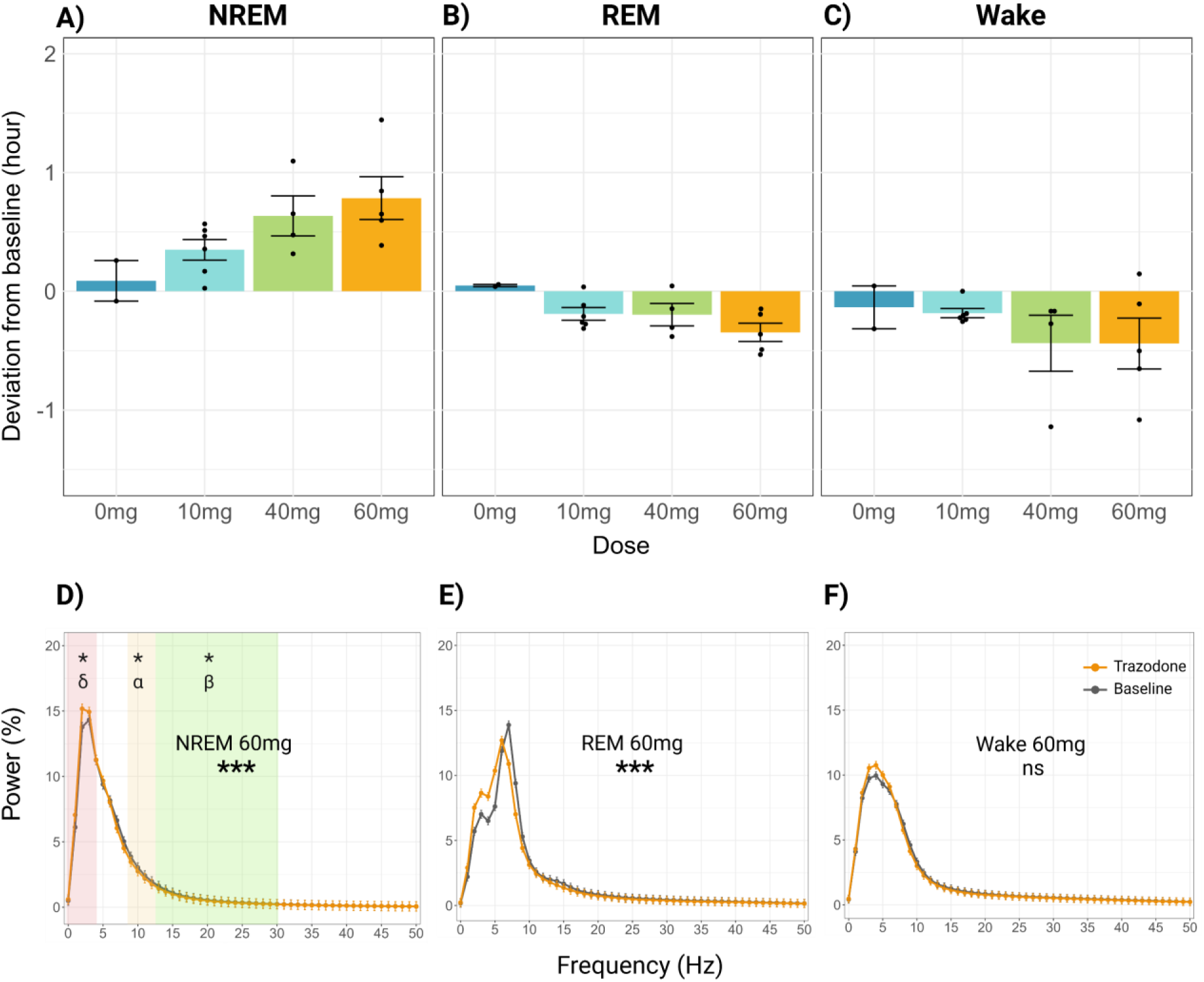
Effects of the highest dose of trazodone (60mg/kg) on NREM sleep and delta power persist after 6 consecutive days of treatment. Changes from baseline in the duration of each vigilance state during the 12-hour light phase, following 6 days of trazodone administration (A-C). The x-axes indicate the trazodone dose. Normalized EEG power spectra on day 6 of trazodone administration at 60mg/kg dose (D-F). Data are expressed as the mean ± SEM. A-C: Mixed-design ANOVA followed by Tukey’s HSD test. D-F: Repeated measures ANOVA followed by Tukey’s HSD test. ^∗^p < 0.05, ^∗∗^p < 0.01, ^∗∗∗^p < 0.001. δ = delta frequency, α = alpha frequency, β = beta frequency.

The effects of trazodone on EEG power spectra remained evident on Day 6 (Fig. 3B) when compared to baseline. Two additional mice were excluded from the power spectra analysis because the EEG from the frontal electrode had high levels of artifact (we were able to reliably use the parietal electrode EEG for the vigilance state scoring). During NREM sleep, a repeated-measures ANOVA revealed a significant interaction between treatment condition (baseline vs. treatment) and frequency at the frontal electrode (F(1.70, 10.21) = 9.01, p = 0.0001) with the Greenhouse-Geisser correction applied. Post hoc Tukey tests indicated a significant increase in delta power compared to baseline (t(4) = 8.02, p = 0.0134), accompanied by reductions in alpha (t(4) = - 7.54, p = 0.0167) and beta (t(4) = - 8.26, p = 0.0120) power. During REM sleep, a significant treatment × frequency interaction was also observed (F(4, 16) = 13.56, p < 0.0001), although post hoc tests did not identify any significant differences. During wake, no significant interaction was detected. The data for the 40mg/kg and 10mg/kg doses also showed significant power spectra shifts and is provided in the Supplementary Materials (see Supplementary Fig. S4).

## Discussion

Trazodone has robust hypnotic effects and is often prescribed clinically for off-label use to treat insomnia; however, preclinical research in mice on the sleep-promoting effects of trazodone remains limited. Here, we validated a novel voluntary oral administration protocol for treating mice with trazodone. Our findings demonstrate that acute trazodone administration increases NREM sleep duration dose-dependently, at the expense of REM sleep and wakefulness. Specifically, a single dose of 60mg/kg produced an average increase of ∼1.5 hour duration of NREM sleep. During NREM sleep, delta power was dose-dependently enhanced, while alpha power was correspondingly reduced. These effects on sleep architecture during the 12-hour rest phase persisted, though to a lesser degree, following six consecutive days of administration. Specifically, on Day 6, the highest dose of trazodone tested (60mg/kg) produced an average increase of ∼45 min duration of NREM sleep, accompanied by an increase in delta power.

Pre-clinical data in mouse models is limited and shows inconsistent results. In a tauopathy mouse model (rTg4510), trazodone did not increase SWS, but did increase REM sleep duration after 8 weeks of treatment [20]. In contrast, similar to our findings, a study in rats showed an increase in the duration of NREM sleep during the 12-hour rest phase following trazodone administration [19]. However, the study did not observe as strong nor persistent effect of trazodone on NREM sleep, presumably due to the lower doses used in their study (2.5 mg/kg and 10mg/kg) [19]. To our knowledge, our study is the first rodent study that successfully induced robust enhancement in SWS, the effect of which persisted for 6 days.

The SWS promoting effects of trazodone have been well reported in clinical studies [4–12]. However, the effects of trazodone on REM sleep have been inconsistent across studies. In our study, we observed a reduction in REM duration, a finding that aligns with the initial human study by Montgomery et al. [4]. Some studies have also reported a slight reduction in REM duration [23] and an increase in REM latency [6,8,11,23]. However, many studies have found no significant change in REM duration [5,7,9,10,12]. The length of the treatment may account for these inconsistencies, as the reduction in REM duration we observed in mice was less pronounced by Day 6 of administration.

The present study has several limitations that warrant further exploration, particularly regarding the duration of the treatment. Evaluating the effects of chronic oral administration of trazodone (beyond 6 days) and its sustained impact on sleep quality, especially SWS, is crucial, as clinical usage typically extends to longer durations and continuous improvement in SWS may be essential for observing tangible therapeutic benefits in the context of neurodegenerative disease. Furthermore, the observed reductions in REM sleep necessitate additional investigation to fully understand the long-term implications of trazodone use. The potential sex differences in the effects of trazodone should also be investigated further.

In summary, we have validated a novel, voluntary oral administration protocol for trazodone in mice. Using this protocol, we have shown that, similarly to humans, trazodone has SWS-enhancing properties in mice, and that the effects are dose-dependent and maintained, although to a lesser degree, after 6 days of treatment. The voluntary oral administration protocol overcomes the previous challenges associated with administering trazodone to mice when studying sleep and will benefit future preclinical research exploring the therapeutic potential of trazodone in mouse models of disease.

## Acknowledgement

This work was funded by Natural Sciences and Engineering Research Council of Canada Discovery Grant (RGPIN/03909-2021, BAK), Vice-President Research Undergraduate Student Research Award (MA), Canada Research Chair (CRC-2020-00047, BAK), and Canada Foundation for Innovation (CFI-41428, BAK). We would like to thank the SFU animal research staff for their support and care for the animals used in this study. We also thank the following research assistants: Katherine Mantel, Emad Shams, and Cody Stevens.

## References

1. Stahl SM. Mechanism of Action of Trazodone: a Multifunctional Drug. CNS Spectr. 2009;14(10):536–546. doi: 10.1017/S1092852900024020.

2. Bossini L, Casolaro I, Koukouna D, Cecchini F, Fagiolini A. Off-label uses of trazodone: a review. Expert Opin Pharmacother. 2012;13(12):1707–1717. doi: 10.1517/14656566.2012.699523.

3. Coin A, Malara A, Noale M, Trevisan C, Devita M, Abbatecola AM, et al. Real-World Use of Trazodone in Older Persons in Long Term Care Setting: A Retrospective Study. Int J Geriatr Psychiatry. 2024;39(11):e70009. doi: 10.1002/GPS.70009.

4. Montgomery I, Oswald I, Morgan K, Adam K. Trazodone enhances sleep in subjective quality but not in objective duration. Br J Clin Pharmacol. 1983;16(2):139. doi: 10.1111/J.1365-2125.1983.TB04977.X.

5. Haffmans PMJ, Vos MS. The effects of trazodone on sleep disturbances induced by brofaromine. European Psychiatry. 1999;14(3):167–171. doi: 10.1016/S0924-9338(99)80736-6.

6. Kaynak H, Kaynak D, Gözükirmizi E, Guilleminault C. The effects of trazodone on sleep in patients treated with stimulant antidepressants. Sleep Med. 2004;5(1):15–20. doi: 10.1016/j.sleep.2003.06.006.

7. Le Bon O, Murphy JR, Staner L, Hoffmann G, Kormoss N, Kentos M, et al. Double-blind, placebo-controlled study of the efficacy of trazodone in alcohol post-withdrawal syndrome: polysomnographic and clinical evaluations. J Clin Psychopharmacol. 2003;23(4):377–383. doi: 10.1097/01.JCP.0000085411.08426.D3.

8. Mouret J, Lemoine P, Minuit MP, Benkelfat C, Renardet M. Effects of trazodone on the sleep of depressed subjects--a polygraphic study. Psychopharmacology (Berl). 1988;95 Suppl(1). doi: 10.1007/BF00172629.

9. Roth AJ, Mccall WV, Liguori A. Cognitive, psychomotor and polysomnographic effects of trazodone in primary insomniacs. J Sleep Res. 2011;20(4):552–558. doi: 10.1111/J.1365-2869.2011.00928.X.

10. Saletu-Zyhlarz GM, Abu-Bakr MH, Anderer P, Gruber G, Mandl M, Strobl R, et al. Insomnia in depression: differences in objective and subjective sleep and awakening quality to normal controls and acute effects of trazodone. Prog Neuropsychopharmacol Biol Psychiatry. 2002;26(2):249–260. doi: 10.1016/S0278-5846(01)00262-7.

11. Scharf MB, Sachais BA. Sleep laboratory evaluation of the effects and efficacy of trazodone in depressed insomniac patients. J Clin Psychiatry. 1990;51 Suppl(9 SUPPL.):13–17.

12. Wang J, Liu S, Zhao C, Han H, Chen X, Tao J, et al. Effects of Trazodone on Sleep Quality and Cognitive Function in Arteriosclerotic Cerebral Small Vessel Disease Comorbid With Chronic Insomnia. Front Psychiatry. 2020;11. doi: 10.3389/FPSYT.2020.00620.

13. Xie L, Kang H, Xu Q, Chen MJ, Liao Y, Thiyagarajan M, et al. Sleep drives metabolite clearance from the adult brain. Science (1979). 2013;342(6156):373–377. doi: 10.1126/SCIENCE.1241224/SUPPL_FILE/XIE-SM.PDF.

14. Fultz NE, Bonmassar G, Setsompop K, Stickgold RA, Rosen BR, Polimeni JR, et al. Coupled electrophysiological, hemodynamic, and cerebrospinal fluid oscillations in human sleep. Science (1979). 2019;366(6465):628–631. doi: 10.1126/SCIENCE.AAX5440/SUPPL_FILE/AAX5440_FULTZ_SM.PDF.

15. Bedward A, Kaur J, Seedat S, Donohue H, Kow CS, Rasheed MK, et al. Pharmacological interventions to improve sleep in people with Alzheimer’s disease: a meta-analysis of randomized controlled trials. Expert Rev Neurother. 2024;24(5):527–539. doi: 10.1080/14737175.2024.2341004.

16. McCleery J, Sharpley AL. Pharmacotherapies for sleep disturbances in dementia. Cochrane Database Syst Rev. 2020;11(11). doi: 10.1002/14651858.CD009178.PUB4.

17. Eyob E, Shaw JS, Bakker A, Munro C, Spira A, Wu M, et al. A Randomized-Controlled Trial Targeting Cognition in Early Alzheimer’s Disease by Improving Sleep with Trazodone (REST). J Alzheimers Dis. 2024;101(s1):S205–S215. doi: 10.3233/JAD-230635.

18. La AL, Walsh CM, Neylan TC, Vossel KA, Yaffe K, Krystal AD, et al. Long-Term Trazodone Use and Cognition: A Potential Therapeutic Role for Slow-Wave Sleep Enhancers. J Alzheimers Dis. 2019;67(3):911–921. doi: 10.3233/JAD-181145.

19. Lelkes Z, Obál F, Alföldi P, Erdos A, Rubicsek G, Benedek G. Effects of acute and chronic treatment with trazodone, an antidepressant, on the sleep-wake activity in rats. Pharmacol Res. 1994;30(2):105–115. doi: 10.1016/1043-6618(94)80002-2.

20. de Oliveira P, Cella C, Locker N, Ravindran KKG, Mendis A, Wafford K, et al. Improved Sleep, Memory, and Cellular Pathological Features of Tauopathy, Including the NLRP3 Inflammasome, after Chronic Administration of Trazodone in rTg4510 Mice. J Neurosci. 2022;42(16):3494–3509. doi: 10.1523/JNEUROSCI.2162-21.2022.

21. Kent BA, Strittmatter SM, Nygaard HB. Sleep and EEG Power Spectral Analysis in Three Transgenic Mouse Models of Alzheimer’s Disease: APP/PS1, 3xTgAD, and Tg2576. J Alzheimers Dis. 2018;64(4):1325–1336. doi: 10.3233/JAD-180260.

22. Kent BA, Michalik M, Marchant EG, Yau KW, Feldman HH, Mistlberger RE, et al. Delayed daily activity and reduced NREM slow-wave power in the APPswe/PS1dE9 mouse model of Alzheimer’s disease. Neurobiol Aging. 2019;78:74–86. doi: 10.1016/J.NEUROBIOLAGING.2019.01.010.

23. van Bemmel AL, Havermans RG, van Diest R. Effects of trazodone on EEG sleep and clinical state in major depression. Psychopharmacology (Berl). 1992;107(4):569–574. doi: 10.1007/BF02245272/METRICS.

