## Supplementary_Material for "The Effects of Acute Trazodone Administration on Sleep in Mice"

### Supplementary Materials

#### Pilot study to test palatable food to mix with trazodone

##### Animals

All experimental protocols were approved by the Simon Fraser University Animal Care and Use Committee (Protocol #1365P-23). A total of n=24 C57BL6/J mice (female n = 11, age 8-13 months) were used to develop the voluntary oral administration protocol. Out of the 24 mice, 18 mice were from Charles River Laboratories (Senneville, Quebec) and 6 mice were bred in-house at the Simon Fraser University facility. Animals were either single- or grouped-housed (up to 3 mice) under a 12:12h light/dark cycle.

##### Method

Trazodone hydrochloride (HCl) powder (catalog number T6154; Sigma-Aldrich) was mixed directly with a palatable food (only for Nutella) or dissolved in distilled water to achieve 10-50mg/ml concentration, and then, mixed with a palatable food. The animals either received food ad libitum or were maintained on a food-restricted diet. All the animals were first habituated to ingest the specific type of treat mixed with a small amount of water (0.8-1.2ml / kg BW). Once they were habituated to ingest the given treat within 20 min, the animals were introduced to the same treat mixed with trazodone.

**Table S1 The type of food tested**

| Type of food | Number of mice that consumed the full amount of the food-trazodone mixture | Number of days the food-trazodone mixture was provided (after habituation) | Tested dose | Food restriction |
|---|---|---|---|---|
| Strawberry-flavored treat (VitaKraft® Drops with Strawberry) | 24/24 | 5.5 | 0, 5, 25, 40, 50, 60 mg/kg BW | Yes |
| Peanut Butter | 1/6 | 2 | 50 mg/kg BW | No |
| Strawberry Jam | 1/5 | 7 | 50 mg/kg BW | Yes |

| | | | | |
|---|---|---|---|---|
| Strawberry Jam + Peanut butter | 0/4 | 4 | 50 mg/kg BW | Yes |
| Laboratory Rodent Diet (Lab Diet 5001) | 0/4 | 1 | 50 mg/kg BW | Yes |
| Strawberry Milkshake (Ensure®) | 0/4 | 2 | 50 mg/kg BW | Yes |
| Froot Loops (Kelloggs) | 0/6 | 2 | 50 mg/kg BW | No |
| Nutella | 1/5 | 6 | 50 mg/kg BW | No |

#### Supplementary Results

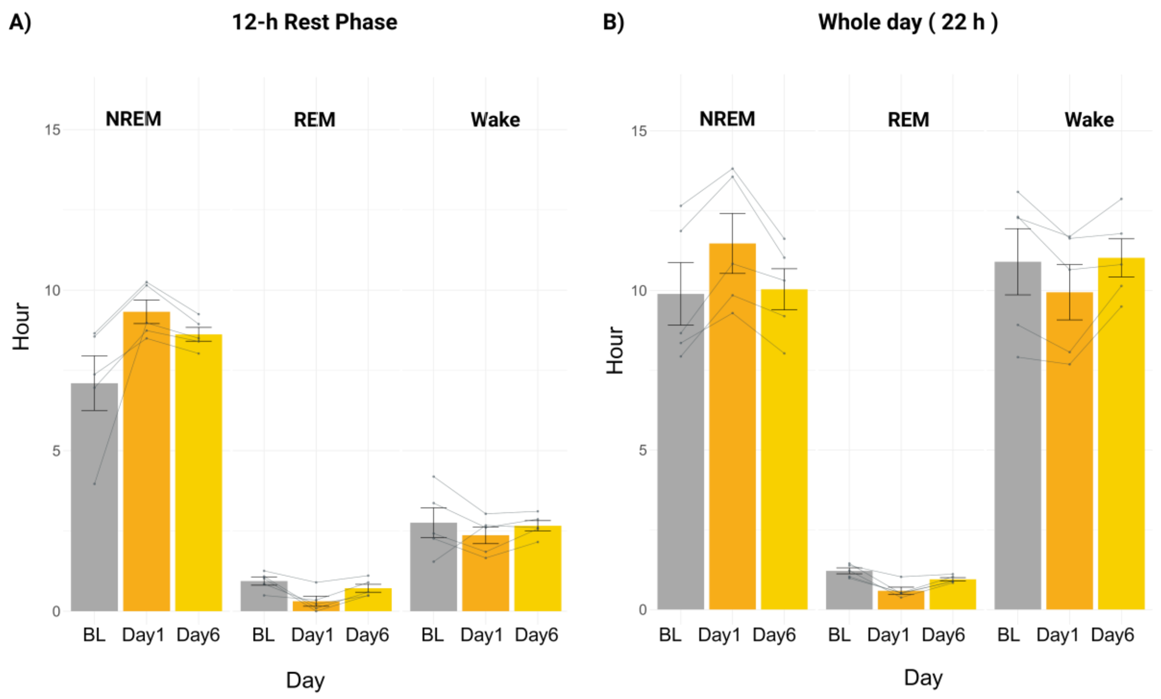


**Supplementary Figure S1**. The difference in NREM, REM, and wake duration between baseline, Day 1, and Day 6 of trazodone administration, expressed in hours along the y-axis, for mice treated with trazodone HCL mixed with palatable food at the 60mg/kg dose (A) during 12-h rest phase and (B) across the whole day (22 hours since recordings were paused during the first and last hour of the 12-h active (dark) phase to minimize noise from feeding standard pellets and administering trazodone, and to re-connect wireless EEG.) Data are expressed as the mean ± SEM.

**A) Day 1**

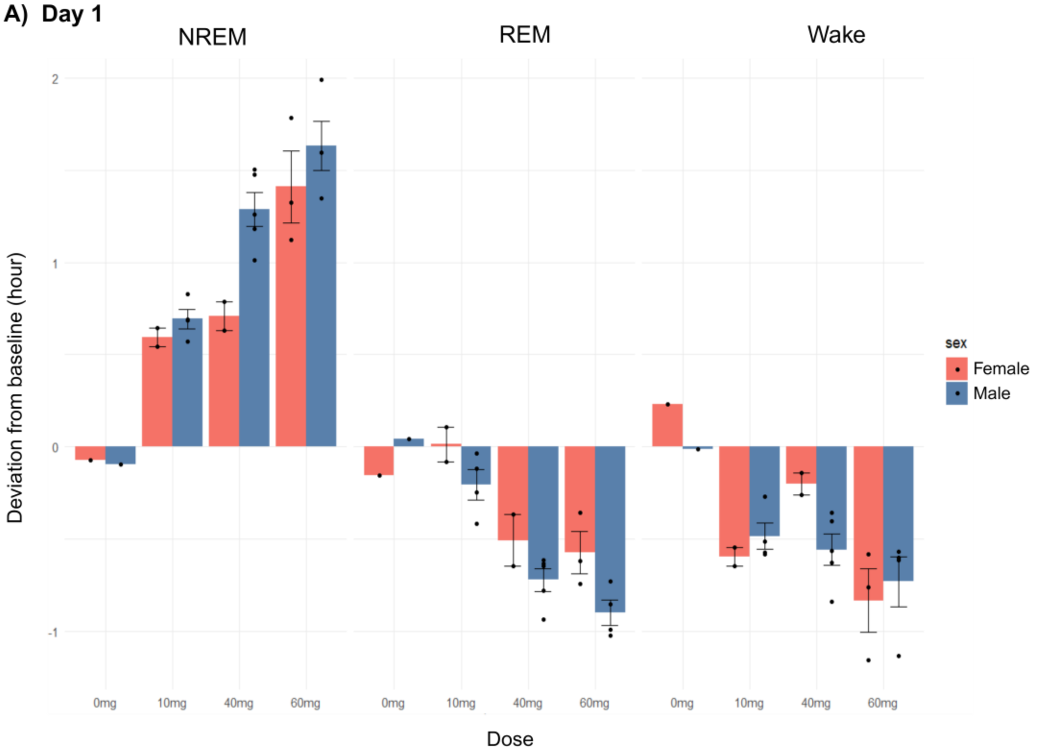
NREM
REM
Wake
Deviation from baseline (hour)
2
1
0
-1
0mg
10mg
40mg
60mg
Dose
sex
Female
Male

**B) Day 6**

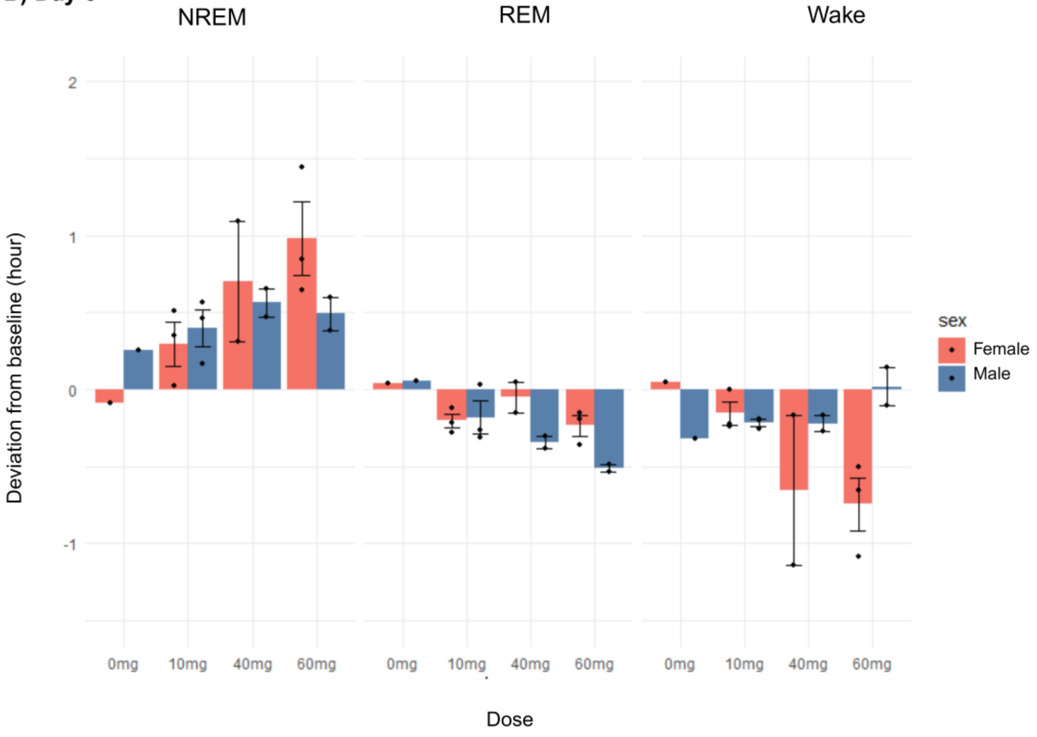
NREM
REM
Wake
Deviation from baseline (hour)
2
1
0
-1
0mg
10mg
40mg
60mg
Dose
sex
Female
Male

**Supplementary Figure S2**. The difference in NREM, REM, and wake duration between trazodone administration day and baseline, expressed in hours along the y-axis, for mice treated with trazodone HCL mixed with palatable food. Female data are shown in red and male data are shown in blue. The x-axes show the dose administered. (A) Day 1 and (B) Day 6 of administration. Data are expressed as the mean ± SEM.

**Day 1 Parietal electrode**

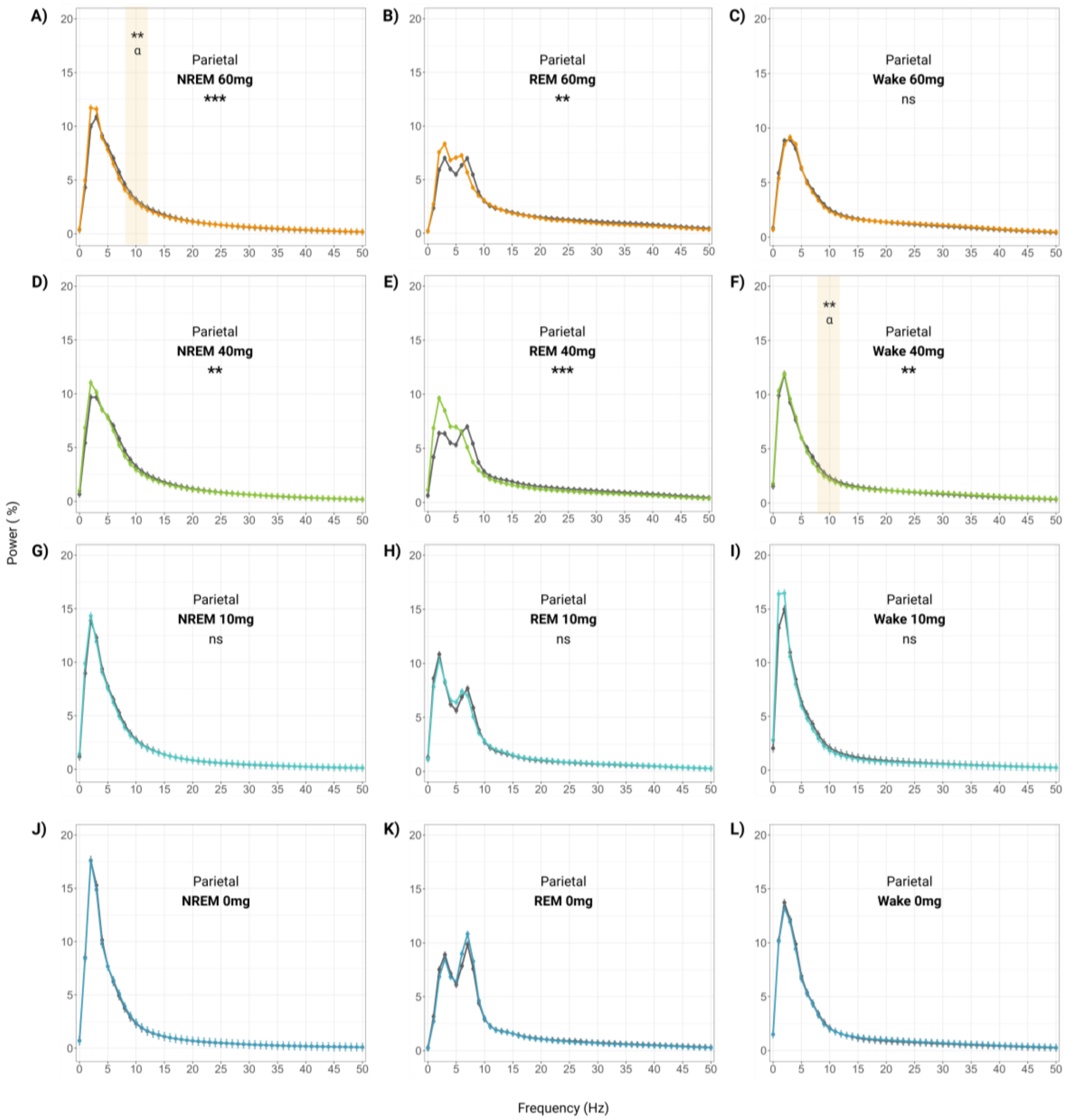


**Supplementary Figure S3.** EEG power spectra recorded by the parietal electrode during NREM, REM, and wake following a single trazodone dose of 60mg/kg (A-C),

40mg/kg (D-F), 10mg/kg (G-I), and 0mg/kg (J-L). Data are expressed as the mean ± SEM. Repeated-measures ANOVA followed by Tukey's HSD test. $* p < 0.05$, $** p < 0.01$, $*** p < 0.001$.

#### Day 6 Frontal electrode

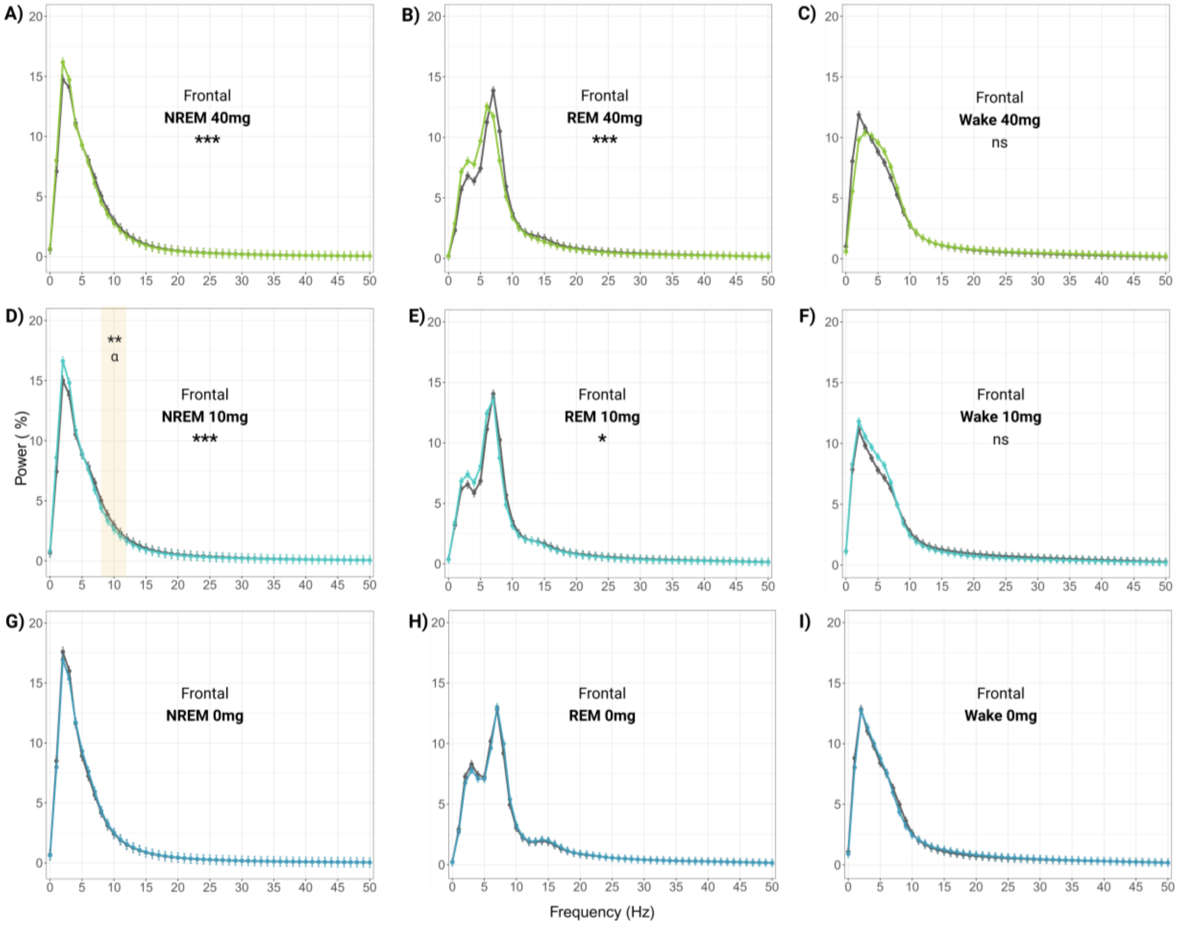


**Supplementary Figure S4**. EEG power spectra recorded by the frontal electrode during NREM, REM, and wake following Day 6 of daily trazodone administration at the dose of 40mg/kg (A-C), 10mg/kg (D-F), and 0mg/kg (G-I). Data are expressed as the mean ± SEM. Repeated-measures ANOVA followed by Tukey's HSD test. $* p < 0.05$, $** p < 0.01$, $*** p < 0.001$.
